# Rab6a enables BICD2/dynein-mediated trafficking of human papillomavirus from the *trans-*Golgi network during virus entry

**DOI:** 10.1101/2024.05.21.595194

**Authors:** Jeongjoon Choi, Kaitlyn Speckhart, Billy Tsai, Daniel DiMaio

## Abstract

Rab GTPases control intracellular vesicular transport, including retrograde trafficking of human papillomavirus (HPV) during cell entry, guiding the virus from the endosome to the *trans-*Golgi network (TGN), the Golgi apparatus, and eventually the nucleus. Rab proteins that act prior to the arrival of HPV to the TGN have been identified, but Rab proteins operating in later stages of entry remain elusive. Here, we report that knockdown of Rab6a impairs HPV entry by impeding intra-Golgi transport of the incoming virus, resulting in HPV accumulation in the TGN. Rab6a supports HPV trafficking by facilitating the association of HPV with dynein, a motor protein complex, and BICD2, a dynein adaptor. L2 can bind directly to GTP-Rab6a *in vitro*, and excess of either GTP-Rab6a or GDP-Rab6 inhibits HPV entry, suggesting that cycling between GDP- Rab6 and GTP-Rab6 is critical. Notably, Rab6a is crucial for HPV-BICD2 and HPV-dynein association in the TGN of infected cells, but the HPV-dynein association in the endosome does not require Rab6a. Our findings reveal an important feature of the molecular basis of intra-Golgi trafficking of HPV and identify potential targets for therapeutic approaches to inhibit HPV infection.

## Introduction

Rab GTPases control intracellular vesicular transport of cellular proteins (1–3). Rab proteins also play critical roles in viral infections, including infection by human papillomaviruses (HPVs) (4–7). HPVs, non-enveloped, double-stranded DNA viruses, are responsible for 5% of human cancer, including essentially all cervical cancer (8). Several Rab proteins are involved in HPV entry (6), and some of which have been assigned specific roles in the entry process. Rab9a and Rab7 act at relatively early steps in this process, prior to the arrival of HPV at the *trans-*Golgi network (TGN) (9–11). Here, we show that Rab6a acts in a relatively late phase of HPV trafficking during virus entry, by coupling HPV to dynein to allow HPV to exit from the TGN.

The HPV virion is composed of 360 molecules of the major capsid protein, L1, and up to 72 molecules of the minor capsid protein, L2 (12). L1 is responsible for HPV binding to the cell surface, while L2 is critical for the trafficking of the viral genome to the nucleus where viral DNA (vDNA) replication and viral gene expression occur (13, 14). After internalization of HPV, viral components including vDNA reside within vesicular retrograde trafficking compartments during the entry process as HPV is transported from endosome to the TGN, through the Golgi apparatus and possibly the endoplasmic reticulum (ER) *en route* to the nucleus (15, 16) (17–19). After initially localizing in endosomes, endosome acidification and γ-secretase activity allow a C-terminal cell-penetrating peptide on the L2 capsid protein to drive a segment of L2 to protrude across the endosomal membrane into the cytosol, so that L2 can interact with various cytoplasmic host factors to enable retrograde trafficking of the incoming virus particle (20, 21). L2 uses this mechanism to associate directly with multiple trafficking factors including retromer (22), sorting nexin (SNX) 17 and 27 (23, 24), dynein (25), retriever (26), and COPI (27). L2 also associates directly or indirectly with Rab9a (11) and Rab7 (10, 11). Most of these factors are involved in relatively early stages of entry, from endosome-to-TGN trafficking of HPV. How HPV traffics from TGN to *cis*-Golgi is not known.

Dynein, a multi-protein complex, plays a pivotal role in orchestrating the intracellular transport of membrane vesicles along microtubules. Dynein associates with L2 during entry, and HPV entry is impaired by chemical inhibition of dynein, mutations in L2 that impair its interaction with dynein, or knockdown of dynein component(s) (25, 28, 29). In addition, RanBP10, a dynein adaptor, facilitates the nuclear import of the HPV vDNA-L2 complex (29). However, beyond this late step in entry (i.e., nuclear import), it remains largely unclear how dynein contributes to the HPV entry during other steps. We recently reported that another dynein adaptor, BICD2, promotes HPV trafficking from endosome to TGN and beyond (30). HPV L2 can bind directly to BICD2 and knockdown of BICD2 inhibits the HPV-dynein interaction and results in HPV accumulation in endosomes and the TGN.

In this study, we show that Rab6a supports HPV type 16 (HPV16) entry by binding to L2 and linking HPV to dynein. Unlike other Rab proteins that associate with HPV in endosomes, Rab6a engages HPV later during entry in the TGN, suggesting that Rab6a plays its role in relatively late stages in the intracellular journey of HPV. Importantly, HPV-BICD2 association and HPV- dynein association at 16 hours post infection require Rab6a, and Rab6a is critical for exit of HPV from the TGN during entry. An excess of either GTP- or GDP-bound forms of Rab6a impairs HPV infection, suggesting that cycling of Rab6a is critical for HPV entry.

## Results

### Rab6a is required for HPV egress from *trans*-Golgi during HPV entry

Although Rab6a is required for infection by HPV16 pseudovirus (PsV) (17), its specific role in HPV entry is unknown. HPV16 PsV consists of a capsid composed of L1 and FLAG-tagged L2 (i.e., L2 with a 3xFLAG tag appended to the C-terminus of L2) capsid containing a reporter plasmid expressing GFP or Gaussia luciferase (Gluc). To investigate the role of Rab6a in HPV entry, we assessed the infectivity of HPV16 PsV containing a plasmid expressing GFP in HeLa S3 cells transfected with <u>n</u>on-targeting <u>c</u>ontrol siRNA (siNC) or siRNA targeting Rab6a expression (siRab6a). Rab6a knockdown was documented by western blotting (Fig. 1A and S1A). Infectivity was measured by flow cytometry for GFP fluorescence at 48 <u>h</u>ours <u>p</u>ost-infection (hpi). This assay measures entry of PsV into the nucleus resulting in expression of the reporter plasmid. Consistent with our previous report (17), Rab6a knockdown significantly inhibited HPV infectivity (∼65-85% reduction compared to control cells) (Fig. 1B and S1B). Another siRNA targeting Rab6a had a similar effect on infectivity (Fig. S1A and S1B). Rab6a depletion also inhibited infection by HPV18 and HPV5 PsVs (Fig. 1C). In addition, Rab6a knockdown inhibited HPV16 PsV infection in human HaCaT skin keratinocytes (Fig. S1C and S1D). Collectively, these data demonstrate the critical role of Rab6a in facilitating the efficient entry of several pathogenic HPV types.

**Fig. 1.**
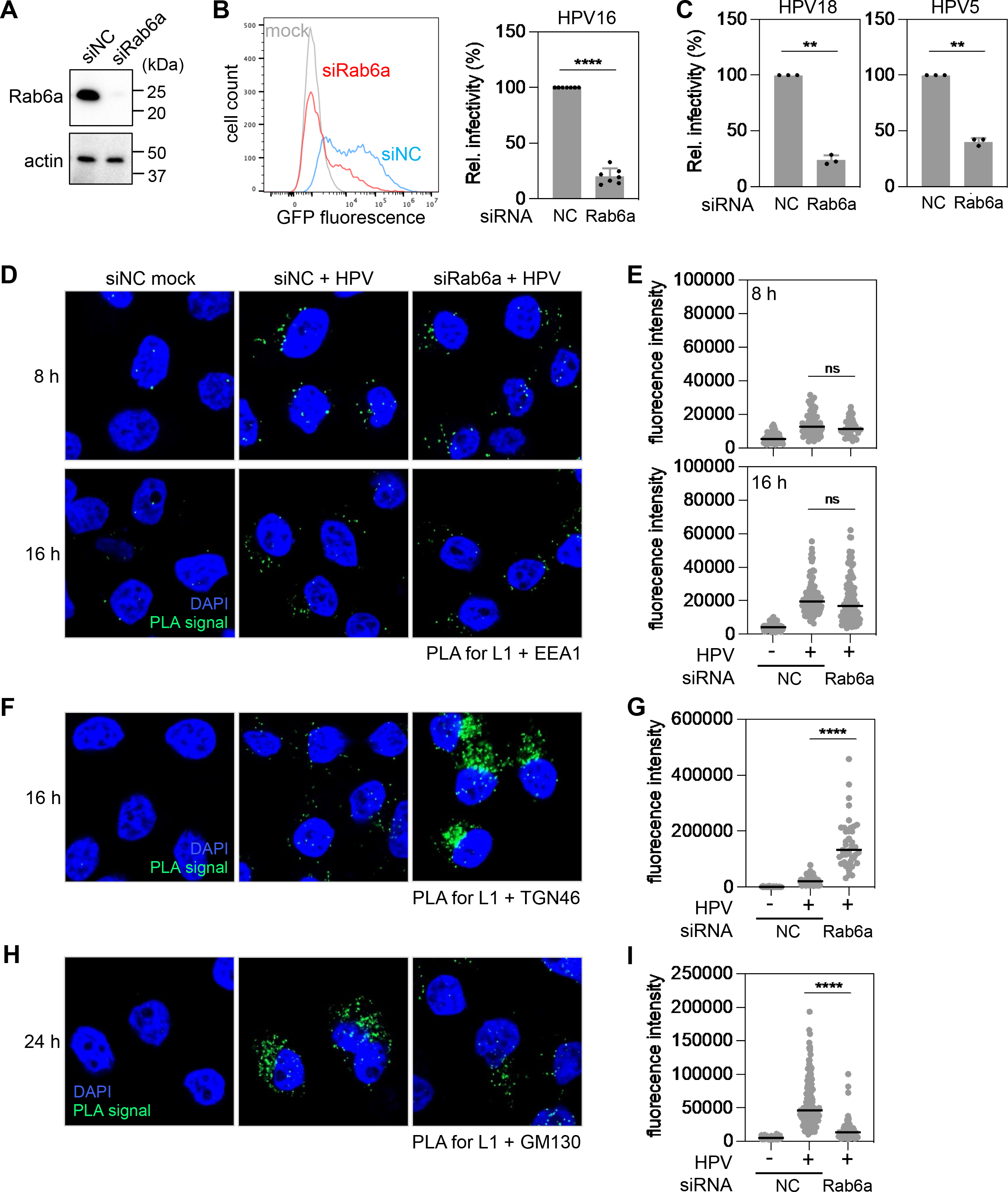
Rab6a knockdown inhibits HPV infection by impairing its intra-Golgi trafficking (from *trans-*Golgi to *cis*-Golgi). (**A**) HeLa S3 cells were transfected with negative control (siNC) or Rab6a-targeting siRNA (siRab6a) and subjected to Western blot analysis using an antibody recognizing Rab6a (top panel) and actin (bottom panel) as a loading control. (**B**) siRNA-treated cells as described in (**A**) were mock-infected or infected at the MOI of ∼2 with HPV16 PsV L2-3XFLAG containing the GFP reporter plasmid. At 48 hpi, GFP fluorescence was determined by flow cytometry. The results are shown as a histogram (*left*) and as percent relative infectivity (based on mean fluorescence intensity) normalized to the siNC treated cells (*right*). Each dot shows the result of an individual experiment. Bars and error bars show mean and standard deviation, respectively. NC, siNC; Rab6a, siRab6a. ****, *P* < 0.0001. (**C**) As in (**B**) except cells were infected with HPV18 and HPV5. (**D**) HeLa S3 cells were transfected with siNC or siRab6a siRNAs and infected with HPV harboring the Gaussia luciferase (Gluc) reporter plasmid at the MOI of ∼200. At 8 and 16 hpi, PLA was performed with antibodies recognizing HPV L1 and EEA1. Mock, uninfected; HPV, infected. PLA signals are green; nuclei are blue (DAPI). Similar results were obtained in two independent experiments. (**E**) The fluorescence of PLA signals was determined from multiple images obtained as in (**D**). Each dot represents an individual cell (*n*>40) and black horizontal lines indicate the mean value of the analyzed population in each group. ****, *P* < 0.0001; ns, not significant. The graph shows results of a representative experiment. (**F**) As in (**D**) except PLA was performed at 16 hpi with antibodies recognizing HPV L1 and TGN46. (**G**) Images as in (**F**) were analyzed as described in (**E**). (**H**) As in (**D**) except PLA was performed at 24 hpi with antibodies recognizing HPV L1 and GM130. (**I**) Images as in (**H**) were analyzed as described in (**E**).

Rab6a is necessary for cellular protein trafficking from Golgi to ER (31–33). To pinpoint the HPV entry step impaired by Rab6a knockdown, we used proximity ligation assay (PLA) to determine HPV localization in HeLa S3 cells infected with HPV16 PsV containing a plasmid expressing Gluc (to avoid interference of GFP fluorescence with the PLA signal). PLA generates a fluorescent signal in intact cells when two proteins recognized by different antibodies are within 40 nm (34). We first examined the proximity of HPV L1 with the endosomal marker protein EEA1. There were negligible L1-EEA1 PLA signals in mock-infected cells. At 8 hpi, similar levels of PLA signals were observed in infected cells transfected with siNC or siRab6a (Fig. 1D and 1E), indicating that Rab6a is not required for HPV to reach the endosome. At 16 hpi, a time when most HPV has largely exited the endosome and entered the TGN in control cells (18, 22), the L1-EEA1 PLA signals were comparable in control and Rab6a knockdown cells (Fig. 1D and 1E), showing that Rab6a is not required for HPV exit from the endosome.

We then conducted PLA for HPV L1 and marker proteins for the TGN (TGN46) or *cis*-Golgi (GM130). At 16 hpi, Rab6a knockdown resulted in much stronger L1-TGN46 PLA signals than in the control cells, indicating that in the absence of Rab6a HPV accumulates in TGN (Fig. 1F and 1G). At 24 hpi, infected Rab6a knockdown cells displayed much lower L1-GM130 PLA signals than control cells (Fig. 1H and 1I), showing that HPV arrival in *cis*-Golgi was impaired in Rab6a depleted cells. The accumulation of HPV in TGN and its depletion from *cis*-Golgi caused by Rab6a knockdown demonstrate that Rab6a facilitates HPV entry by supporting HPV trafficking from the TGN to the *cis*-Golgi.

### Rab6a is in close proximity to HPV at late times post-infection

If Rab6a acts directly on HPV during entry, we predict that Rab6a and HPV will be in close proximity when the virus is in the TGN. To determine is this is the case, we next conducted PLA for L1 and Rab6a at 8 and 16 hpi (Fig. 2A and 2B). Minimal PLA signals were detected in uninfected cells or in cells 8 hpi. In contrast, at 16 hpi, strong L1-Rab6a PLA signals were detected, indicating that HPV is in close proximity to Rab6a during the relatively late stage of entry at which it appears to support trafficking. PLA similarly demonstrated that L2 and Rab6a were in close proximity at 16 hpi (Fig. 2C and 2D). Thus, HPV is in proximity to Rab6a at 16 hpi, but not at 8 hpi (Fig. 2A-2D), consistent with a role of Rab6a in HPV trafficking out of the TGN.

**Fig. 2.**
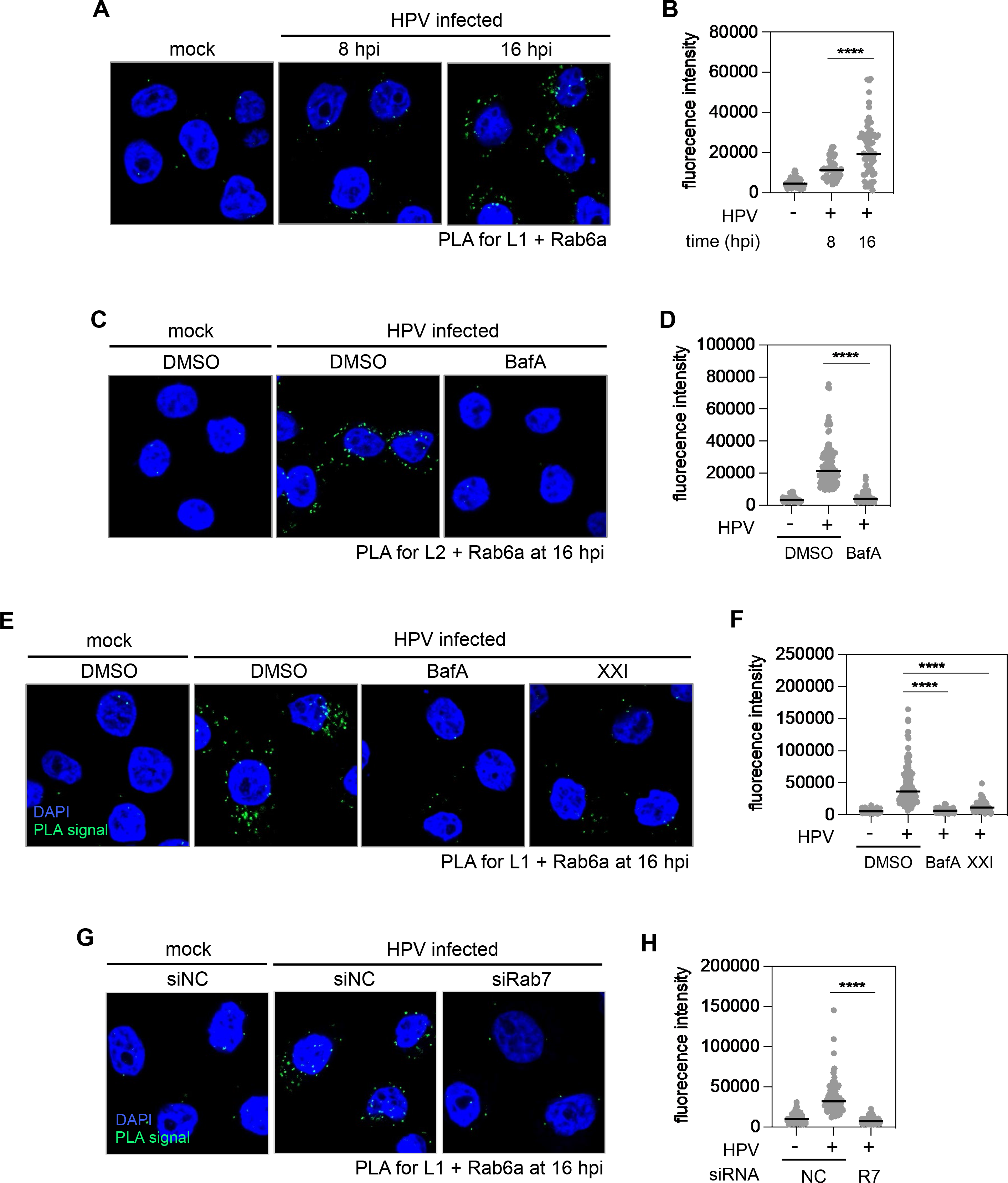
Rab6a engages with HPV at relatively late times during entry. (**A**) HeLa S3 cells were mock-infected or infected at the MOI of ∼200 with HPV16 PsV L2-3XFLAG containing the Gluc reporter plasmid. At 8 and 16 hpi, PLA was performed with antibodies recognizing HPV L1 and Rab6a. PLA signals are green; nuclei are blue (DAPI). Similar results were obtained in two independent experiments. (**B**)The fluorescence of PLA signals was determined from multiple images obtained as in (**A**). Each dot represents an individual cell (*n*>40) and black horizontal lines indicate the mean value of the analyzed population in each group. ****, *P* < 0.0001. The graph shows results of a representative experiment. (**C**) As in (**A**) except DMSO or Bafilomycin A1 (BafA) were added to the medium 30 min prior to infection, and PLA was performed at 16 hpi with antibodies recognizing FLAG (HPV L2) and Rab6a. (**D**) Images as in (**C**) were analyzed as described in (**B**). (**E**) As in (A) except DMSO, BafA, or γ-secretase inhibitor XXI were added to the medium 30 min prior to infection, and PLA was performed at 16 hpi (**F**) Images as in (**E**) were analyzed as described in (**B**). (**G**) HeLa S3 cells were transfected with negative control (siNC) or Rab7-targeting siRNA (siRab7) and infected at the MOI of ∼200 with HPV16 PsV L2-3XFLAG containing the Gluc reporter plasmid. At 16 hpi, PLA was performed with antibodies recognizing HPV L1 and Rab6a. Similar results were obtained in two independent experiments. (**H**) Images as in (**G**) were analyzed as described in (**B**).

Because Rab6a associates with HPV at later stages of entry and is required for exit of HPV from the TGN, we predicted that blocking transport of HPV to the TGN would prevent the HPV- Rab6a interaction. To test this, we used PLA at 16 hpi to assess the association of HPV and Rab6a in cells where HPV trafficking out of the endosome was blocked. Endosomal acidification, γ-secretase activity, and Rab7 are required for HPV trafficking from endosome to TGN at early stages of infection (10, 20, 35, 36). Rab6a association with L1 was markedly reduced by treatment with Bafilomycin A1 (BafA, an inhibitor of endosome acidification), XXI (an inhibitor of γ-secretase), or by Rab7 knockdown (Fig. 2E-2H). Similarly, L2-Rab6a PLA signal at 16 hpi was abolished by BafA (Fig. 2C and 2D). Collectively, these findings show that endosome acidification and the action of Rab7 and γ-secretase are required for HPV to come into proximity with Rab6a. We conclude that arrival of HPV in the TGN is necessary for HPV-Rab6a association.

### HPV-BICD2-dynein interaction in the TGN requires Rab6a

Dynein is known to interact with Rab6a and is thought to play a role in retrograde trafficking of HPV during entry (25, 28, 37, 38). We used PLA at 8 and 16 hpi for L1 and dynein to test whether Rab6a is required for HPV-dynein association in infected cells. In control cells, clear L1-dynein PLA signals were detected at both 8 and 16 hpi, but were negligible in mock-infected cells, indicating that HPV associates with dynein during entry, consistent with previous reports (25, 28, 29). Notably, HPV-dynein interaction was markedly decreased in cells depleted of Rab6a at 16 hpi, but not at 8 hpi (Fig. 3A and 3B), consistent with the association of Rab6a with HPV at 16 hpi, but not 8 hpi (Fig. 2A and 2B). These results indicate that Rab6a plays a critical role in HPV-dynein association at 16 hpi when the virus is in the TGN, but it is not required for HPV-dynein interaction at 8 hpi when the virus is primarily in the endosome.

**Fig. 3.**
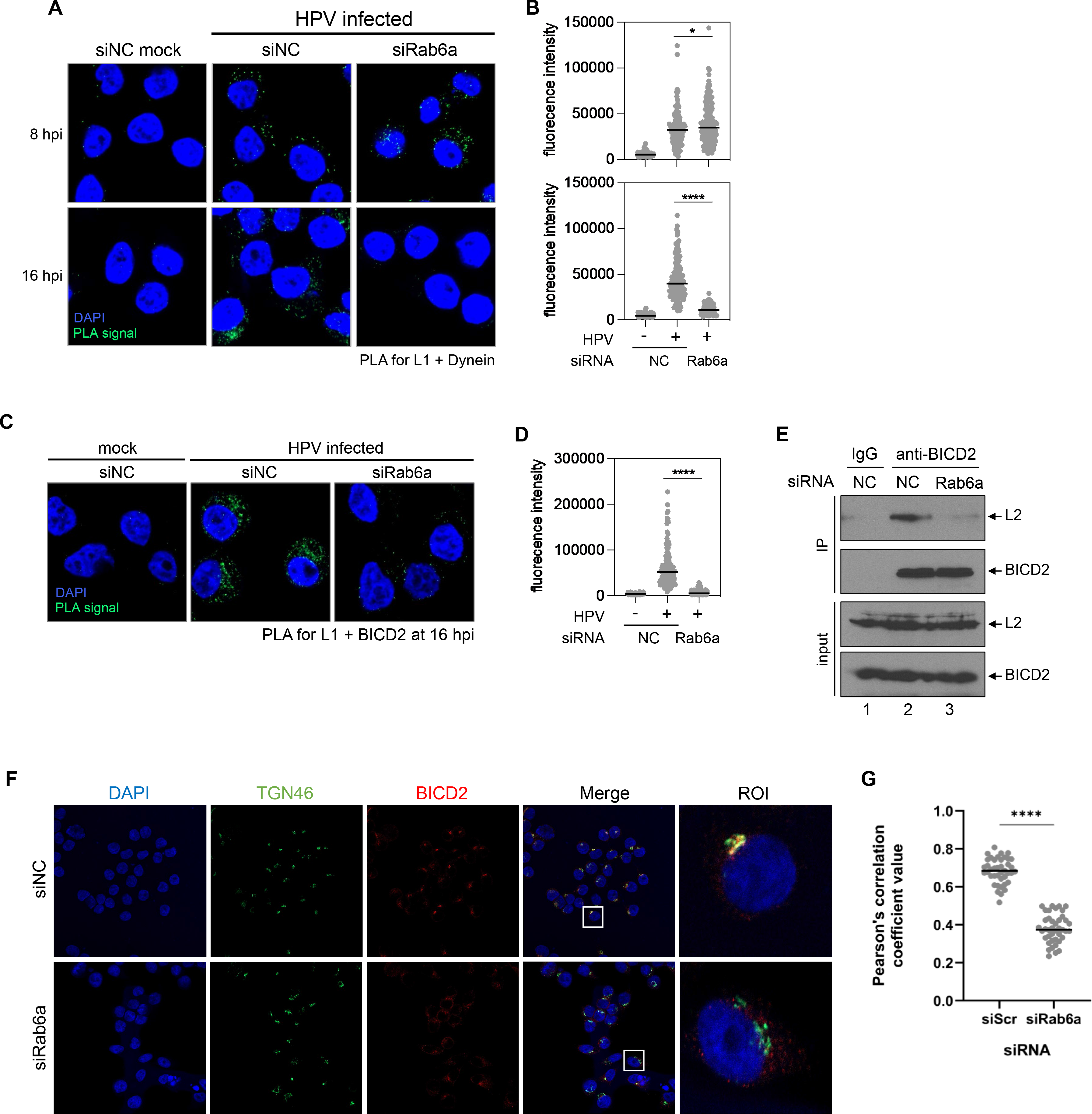
Rab6a is required for HPV-BICD2-dynein association. (**A**) HeLa S3 cells were transfected with siNC or siRab6a siRNAs and infected with HPV harboring the Gluc reporter plasmid at the MOI of ∼200. At 8 and 16 hpi, PLA was performed with antibodies recognizing HPV L1 and dynein. Mock, uninfected; HPV, infected. PLA signals are green; nuclei are blue (DAPI). Similar results were obtained in two independent experiments. (**B**) The fluorescence of PLA signals was determined from multiple images obtained as in (**A**). Each dot represents an individual cell (*n*>40) and black horizontal lines indicate the mean value of the analyzed population in each group. ****, *P* < 0.0001. The graph shows results of a representative experiment. (**C**) As in (**A**) except PLA was performed with antibodies recognizing HPV L1 and BICD2. (**D**) Images as in (**C**) were analyzed as described in (**B**). (**E**) siRNA-transfected cells were infected with HPV harboring the GFP reporter plasmid at the MOI of ∼1. At 16 hpi, cells were lysed and immunoprecipitated with control antibody or antibody recognizing BICD2. Precipitated samples were subjected to Western blot analysis using an antibody recognizing FLAG (HPV L2) and BICD2. (**F**) HeLa S3 cells were transfected with siNC or siRab6a. After 48 h, BICD2 and TGN46 were stained using antibodies recognizing BICD2 and TGN46. Immunofluorescence (IF) images were shown; TGN46, green; BICD2, red; nuclei (DAPI), blue. Merged image shows TGN46 and BICD2 with overlap colored yellow. Similar results were obtained in three independent experiments. (**G**) Pearson’s correlation coefficient values for TGN46 and BICD2 colocalization in multiple images obtained as in (**F**). Each dot represents an individual cell (n>40), and horizontal lines indicate the mean value of the analyzed population in each group. ****, *P* < 0.0001. The graph shows results of a representative experiment.

We previously reported that BICD2, a cargo adaptor for dynein, binds HPV L2 and is required for HPV-dynein association during HPV entry (30, 39, 40). Furthermore, it is well-established that Rab6a interacts with BICD2 and promotes dynein-mediated cargo trafficking (41–44). Thus, we hypothesized that Rab6a may enable the HPV-BICD2 interaction in the TGN and thereby promote HPV-dynein association. We used PLA for L1 and BICD2 at 16 hpi to test whether Rab6a knockdown affects HPV interaction with BICD2. At this time point, L1-BICD2 PLA signals were greatly reduced by Rab6a knockdown (Fig. 3C and 3D), although Rab6a knockdown did not affect expression of BICD2 (Fig. 3E). These results indicate that Rab6a is required for HPV-BICD2 interaction in the TGN at 16 hpi. We note that BICD2 localization at the TGN was reduced by Rab6a KD in uninfected cells (Fig. 3F and 3G). Mislocalization of BICD2 in response to Rab6a KD presumably explains at least in part the decreased HPV-BICD2 association in Rab6a-depleted cells.

As an independent approach to assess the interaction of HPV and BICD2, we performed immunoprecipitation experiments from detergent lysates of control and Rab6a knockdown cells at 16 hpi. The anti-BICD2 antibody, but not the control IgG, co-immunoprecipitated (co-IPed) the L2 protein from control infected cells (Fig. 3E, lanes 1 and 2) as expected, but the amount of L2 protein co-IPed with BICD2 was markedly reduced from lysates of Rab6a knockdown cells (Fig. 3E, lanes 2 and 3). Taken together with the PLA experiments described above, these results indicate that Rab6a is required for optimal HPV-BICD2 interaction in intact cells and cell extracts at 16 hpi and imply that Rab6a facilitates HPV-BICD2-dynein association at the TGN but not at earlier steps in entry.

### Excess GTP- or GDP-bound Rab6a impairs HPV entry

Because the GTP-bound form of Rab proteins typically promotes trafficking of cellular protein cargo (1), we investigated whether GTP-Rab6a promotes HPV entry. We introduced constitutively active (CA) HA-tagged Rab6a mutant (Q72L) into 293TT cells to induce a higher GTP-Rab6a to GDP-Rab6a ratio compared to normal cells (Fig. 4A and 4B). Forty-eight h after transfection with a plasmid expressing CA Rab6a, cells were infected with HPV16 PsV. At 48 hpi, we used flow cytometry to measure GFP fluorescence to determine HPV infection and anti-HA staining to measure expression of the mutant Rab6a protein. This experimental design allowed us to compare HPV entry in the presence or absence of CA Rab6a (i.e., in HA-positive and HA-negative cells, respectively) within the same cell population (Fig. 4C). Contrary to our expectation, cells with a presumably increased GTP-Rab6a to GDP-Rab6a ratio due to the expression of CA Rab6a (Fig. 4B) showed reduced HPV infection compared to cells lacking CA Rab6a (Fig. 4C, left). This result indicates that excess GTP-Rab6a impairs HPV infection.

**Fig. 4.**
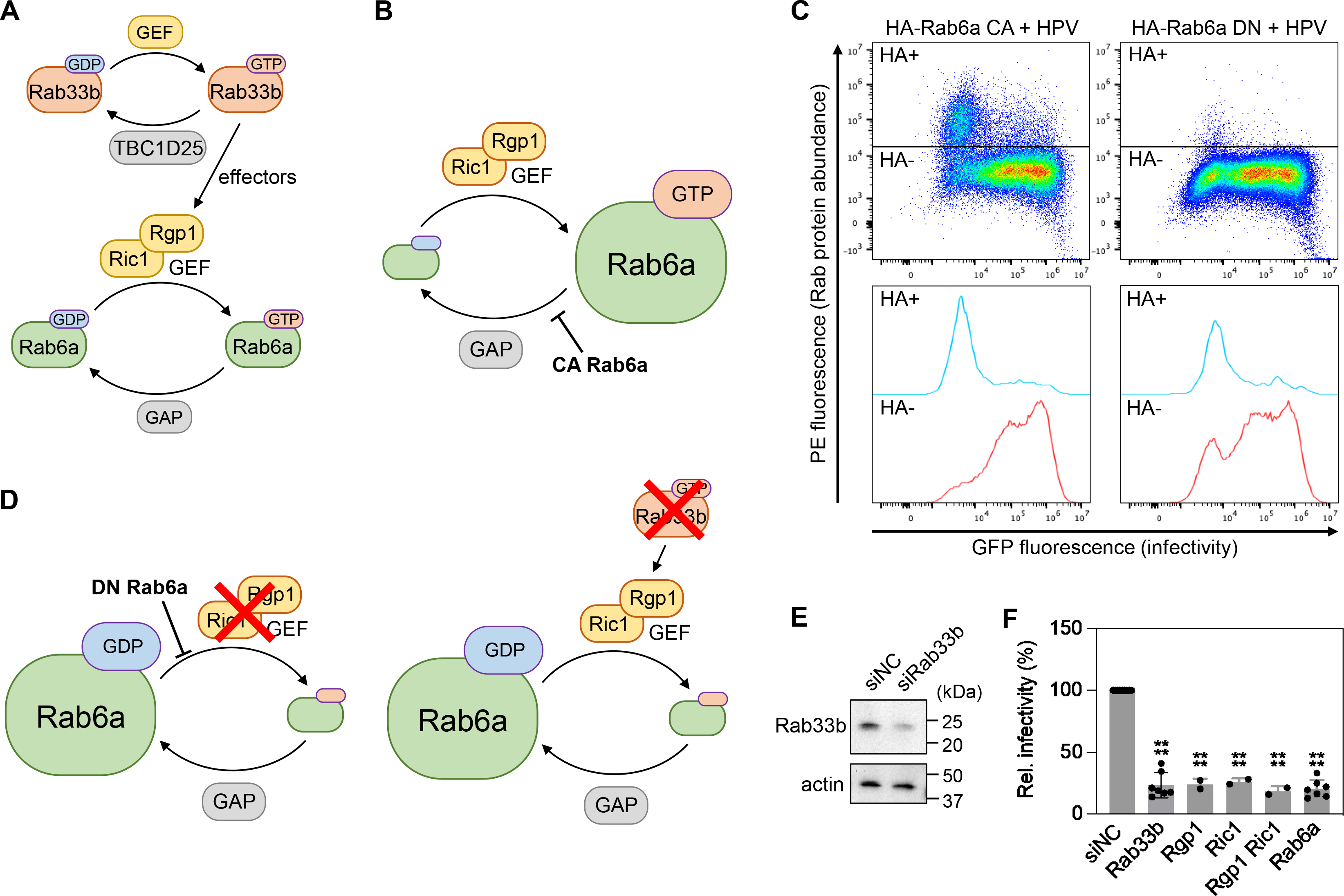
Excess of either GTP-Rab6a or GDP-Rab6a impairs HPV entry. (**A**) Schematic of Rab6a cycling between GTP-bound and GDP-bound forms. Ric1/Rgp1, a Rab6a GEF complex, converts GDP-Rab6a to GTP-Rab6a by exchanging GDP to GTP. A GTPase-activating protein (GAP) hydrolyzing GTP-Rab6a is yet to be identified. Ric1 and Rgp1 are effectors of Rab33b. TBC1D25 is a GAP for Rab33b and a GEP for Rab33b has not been identified. (**B**) Constitutively active (CA) Rab6a mutant is locked in GTP-bound form; thus, cells expressing CA Rab6a accumulate GTP-Rab6a. (**C**) Dominant negative (DN) Rab6a mutant is locked in GDP-bound form; thus, cells expressing DN Rab6a accumulate GDP-Rab6a. Knockdown of Rab33b also results in accumulating GDP-Rab6a. (**D**) 293TT cells were transfected with a plasmid expressing HA-tagged CA or DN Rab6a. At 48 h after transfection, cells were infected at the MOI of ∼2 with HPV16 PsV L2-3XFLAG containing the GFP reporter plasmid. At 48 hpi, samples were stained using PE-conjugated antibody recognizing HA, and flow cytometry was used to determine PE and GFP fluorescence. Cells expressing HA-tagged CA or DN Rab6a were marked as HA+, and those not expressing such proteins were marked as HA-. Representative dot blots are shown in top panels. On the bottom, corresponding histograms are shown. (**E**) HeLa S3 cells were transfected with negative control (siNC) or Rab33b-targeting siRNA (siRab33b) and subjected to Western blot analysis using an antibody recognizing Rab33b (top panel) and actin (bottom panel) as a loading control. (**F**) siRNA-treated cells as described in (**E**), except targeting siRNAs were used as indicated, were infected at the MOI of ∼2 with HPV16 PsV L2-3XFLAG containing the GFP reporter plasmid. At 48 hpi, GFP fluorescence was determined by flow cytometry. Each dot shows the result of an individual experiment. Bars and error bars show mean and standard deviation, respectively. ****, *P* < 0.0001.

Next, we investigated the effect of excess GDP-Rab6a by transiently expressing the dominant negative (DN) Rab6a mutant (T27N), which is predicted to cause accumulation of GDP-bound Rab6a (Fig. 4A and 4D, left). The increased GDP-Rab6a to GTP-Rab6a ratio in cells expressing HA-tagged DN Rab6a also inhibited HPV infection compared to cells lacking DN Rab6a (Fig. 4C, right), indicating that excess GDP-Rab6a impairs HPV infection as well. As an alternative approach to generate cells with excess GDP-Rab6a, we knocked down Rgp1 and Ric1 (Fig. 4D, left), which form a complex that displays guanine nucleotide exchange factor (GEF) activity for Rab6a. Thus, Rgp1/Ric1 KD is predicted to result in increased GDP-Rab6 (Fig. 4D, left). Individual knockdown of Rgp1 or Ric1, as well as combined knockdown of both proteins reduced HPV infection (Fig. 4F), supporting the conclusion that excess GDP-Rab6a impairs HPV infection. Finally, we knocked down Rab33b (Fig. 4E), which recruits GEF to Rab6a (Fig. 4D, right) (45), and tested infectivity and localization of the incoming HPV. Rab33b knockdown, which also is predicted to increase GDP-Rab6a, also inhibited HPV infection (Fig. 4F) and resulted in HPV accumulation at TGN (Fig. S2A and S2B). Taken together, these results show that excess of either GTP-bound or GDP-bound Rab6a impairs HPV entry.

### A C-terminal segment of HPV L2 preferentially interacts with GTP-bound Rab6a *in vitro*

To determine if Rab6a can bind directly to L2 and to identify which portion of L2 interacts with Rab6a, we conducted *in vitro* pull-down experiments using purified HA-tagged CA Rab6a protein (Fig. 5A) and biotinylated peptides corresponding to amino acids 12-44 (designed L2 N in the figure), 299-312 (L2 M), or 434-461 of L2 (L2 C) (Fig. 5B and 5C). Each L2 peptide or 3xFLAG peptide as negative control was incubated with purified CA Rab6a protein, then the peptides were pulled down using streptavidin beads. Of these four different peptides, only peptide 434-461 pulled down CA Rab6a protein (Fig. 5D). This result indicates that the C-terminus portion of L2 (434–461) can interact directly with Rab6a.

**Fig. 5.**
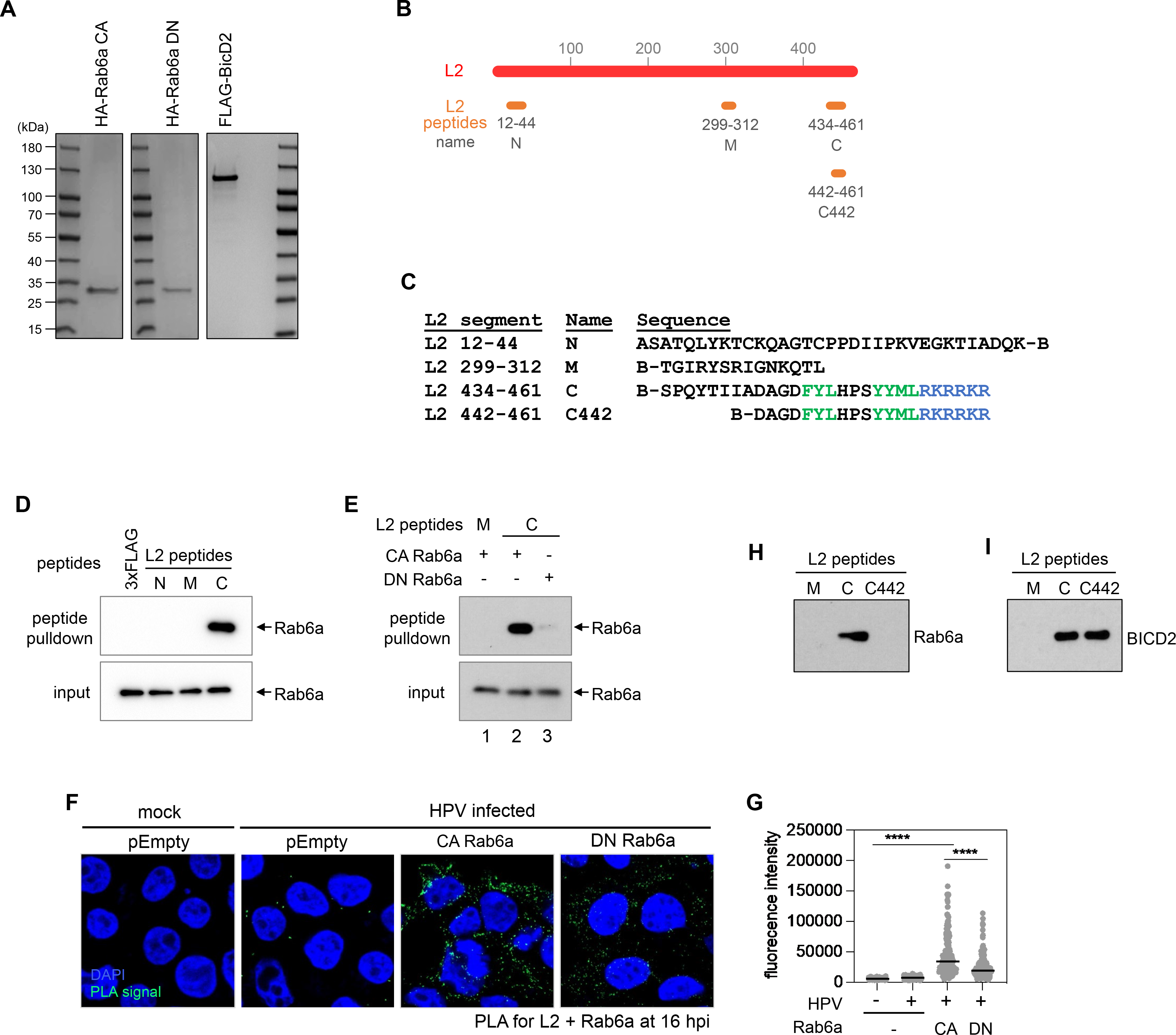
C-terminus segment of L2 interacts with Rab6a, preferentially in its GTP-bound form. (**A**) Purified HA-tagged CA and DN Rab6a and FLAG-tagged BICD2 proteins were separated in SDS-PAGE and visualized by Coomassie staining. (**B**) Schematic of L2 (red) and L2 peptides covers different segments in L2 (orange). Numbers under each peptide indicate amino acid positions from the full-length L2. Name of each peptide was indicated under those numbers. (**C**) Sequences of the HPV16 L2 peptides. B indicates biotin. Amino acids of the retromer binding site are shown in green and those of the CPP are shown in blue. (**D**) Purified HA-tagged CA Rab6a proteins were incubated with indicated biotinylated L2 peptides. Peptide- associated proteins were then pulled down using streptavidin beads and subjected to Western blot analysis using an antibody recognizing HA. Similar results were obtained in two independent experiments. A representative image is shown. (**E**) As in (**D**) except using both CA and DN Rab6a proteins. (**F**) 293TT cells were transfected with a plasmid expressing HA-tagged CA or DN Rab6a, or empty vector (pEmpty). At 48 h after transfection, cells were infected at the MOI of ∼200 with HPV16 PsV L2-3XFLAG containing the Gluc reporter plasmid. At 16 hpi, PLA was performed with antibodies recognizing FLAG (HPV L2) and Rab6a. PLA signals are green; nuclei are blue (DAPI). Similar results were obtained in two independent experiments. (**G**) The fluorescence of PLA signals was determined from multiple images obtained as in (**F**). Each dot represents an individual cell (*n*>40) and black horizontal lines indicate the mean value of the analyzed population in each group. ****, *P* < 0.0001. The graph shows results of a representative experiment. (**H** and **I**) Purified HA-tagged CA Rab6a (**H**) or FLAG-tagged BICD2 (**I**) were incubated with indicated biotinylated L2 peptide (M, C, or C442). Peptide- associated proteins were then pulled down using streptavidin beads and subjected to Western blot analysis using an antibody recognizing FLAG or HA. Similar results were obtained in two independent experiments. Representative images are shown.

We also purified HA-tagged DN Rab6a protein (Fig. 5A) and performed pull-down experiments using the C-terminal L2 peptide and CA or DN Rab6a. As expected, peptide 434- 461 pulled down CA Rab6a whereas peptide 299-312 peptide did not (Fig. 5E, lanes 1 and 2). However, peptide 434-461 pulled down very little DN Rab6a (Fig. 5E, lanes 2 and 3), indicating that L2 preferentially binds to GTP-Rab6a *in vitro* compared to GDP-Rab6a. Consistent with this result with purified components, L2-Rab6a PLA showed higher PLA signals in cells expressing CA Rab6a than in cells expressing DN Rab6a (Fig. 5F and 5G). Taken together, these results show that the HPV L2 C-terminus region (434–461) preferentially interacts with GTP-Rab6a during HPV entry.

### Overlapping segments of HPV L2 are required for *in vitro* interaction with Rab6a and BICD2

Although BICD2 association with HPV requires Rab6a in infected cells (Fig. 3C and 3D), we previously reported *in vitro* binding experiments showing that BICD2 can bind directly to L2 in the absence of Rab6a (30). Here, we used the L2 C peptide to examine if Rab6a and BICD2 bind to the same segment of L2 *in vitro*. When tested individually, purified FLAG-tagged BICD2 and HA-tagged CA Rab6a were both pulled down by peptide 434-461 but not by the control peptide 299-312 (Fig. 5H and 5I). Thus, in this *in vitro* reaction, BICD2 was able to bind this L2 segment in the absence of Rab6a, consistent with our previous report.

We also tested L2-protein binding using a shorter peptide L2 442-461 (designated C442) (Fig. 5B and 5C). Interestingly, this 20-amino-acid-long peptide precipitated as much BICD2 as peptide 434-461, as reported previously (30), but it did not pull down Rab6a (Fig. 5H and 5I).

This result suggests that, although BICD2 and Rab6a both bind to C-terminal segment of L2 in vitro, amino acids 434 to 441 are required for Rab6a binding but not for BICD2 binding.

## Discussion

Rab GTPases play pivotal roles in HPV entry (1–3, 6). In the early stages of entry, Rab7 and Rab9a associate with HPV and regulate the association and dissociation of retromer, thereby enabling endosome-to-TGN retrograde HPV transport (10, 11). This study reveals that upon arrival at the TGN, Rab6a associates with HPV and promotes the assembly or maintenance of an HPV-BICD2-dynein complex, thereby allowing dynein-mediated intra-Golgi HPV trafficking (Fig. 6). This model is based on several findings reported here: (1) Rab6a knockdown inhibits exit of incoming HPV from the TGN and depletes it form the *cis*-Golgi, thus inhibiting infection, supporting the model that HPV exit from the TGN and arrival at the Golgi stacks is important for infection. (2) Rab6a directly binds L2 in vitro and is in close proximity to HPV when it is localized to the TGN at 16 hpi during entry, and (3) Rab6a knockdown inhibits the association of HPV with dynein and its adaptor, BICD2, at 16 hpi, but not at 8 hpi. In addition, COPI/L2 binding also plays a role in HPV transit through the TGN/Golgi (27). Thus, HPV entry is mediated by an intricate retrograde trafficking mechanism that sequentially employs multiple different host factors.

**Fig. 6.**
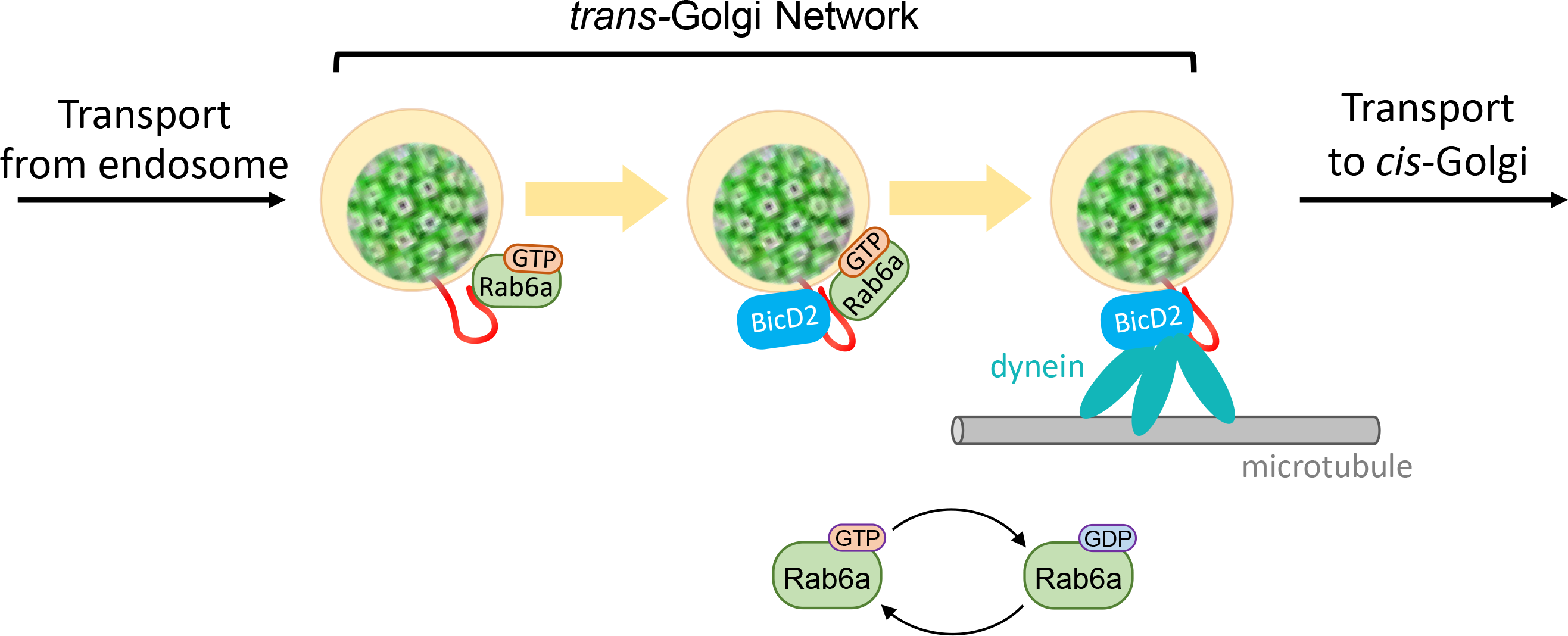
Rab6a promotes HPV trafficking from the trans-Golgi network to cis-Golgi by enabling BICD2/dynein association with HPV during virus entry. After retrograde trafficking of internalized HPV to *trans-*Golgi, L2 associates with GTP-Rab6a, which recruits BICD2, an adaptor for dynein complex. Hydrolysis of GTP-Rab6a promotes dissociation of Rab6a from L2, leaving L2 associated with BICD2/dynein, allowing HPV cargo trafficking on the microtubule to *cis*-Golgi.

L2 directly interacts with Rab6a (Fig. 5), and Rab6a depletion abrogates HPV-BICD2 and HPV-dynein association in HPV-infected cells as assessed by PLA and co-IP at 16 hpi (Fig. 3). Thus, HPV-BICD2 association in the TGN in infected cells requires Rab6a, consistent with the primary localization of Rab6a to the TGN (46, 47). Our results also suggest that the L2-BICD2 and L2-dynein association previously detected by co-IP from cell lysates is likely to be mediated by endogenous Rab6a in the lysates (25, 28, 30).

Rab proteins exist in two forms: an active GTP-bound state necessary for interacting with effector proteins that supports the trafficking of cellular cargo and an inactive GDP-bound state (1, 3). Consistent with this paradigm, L2 preferentially associates with GTP-Rab6a compared to GDP-Rab6a. Typically, excess GTP-Rab6a allows the transport of cellular cargo such as glycosylation enzymes or Shiga toxin, while excess GDP-Rab6a blocks their transport (31, 32). Unlike typical cellular cargo trafficking, however, excess of either the GTP- or the GDP-bound form of Rab6a impairs HPV entry, suggesting that HPV utilizes Rab6a in a different way than does the host cell. Furthermore, Rab6a promotes retrograde trafficking of cellular cargo from the *cis*-Golgi to the ER (31, 32), whereas Rab6a supports retrograde HPV trafficking from the TGN to the *cis*-Golgi, suggesting that Rab6a mediates a different step of retrograde trafficking for HPV compared to cellular protein cargo. Thus, the viral cargo may in effect reprogram the cellular trafficking machinery to support virus entry. We note that Rab6a also regulates anterograde intra-Golgi trafficking of cellular cargo (48–50).

The findings reported here, together with our previously published work, reveal the role of three Rab proteins during HPV entry: Rab6a, Rab7, and Rab9a. The cycling of Rab6a and Rab7 appears to be critical for HPV entry because excess of either GTP- or GDP-bound forms of these proteins inhibits entry (10, 11). In contrast, excess GDP-Rab9a promotes HPV entry, while GTP- Rab9a inhibits it (11). Our results also show that Rab33b is involved in HPV entry, apparently by regulating Rab6a activity. In addition, Rab5 is also important for HPV entry as CA or DN Rab5 impaired infection (9, 46).

Rab proteins function at various stages during HPV entry. Rab5 likely functions at a very early step of HPV entry as it is required for HPV localization to endosome, but it is not known if Rab5 acts on HPV itself or on a cellular protein required for HPV entry into the endosome (9).

Once HPV is in endosomes, endosome acidification and γ-secretase action trigger HPV L2 protrusion across the endosomal membrane, thereby enabling L2 to interact sequentially with Rab9a and later Rab7, which regulate HPV-retromer association. Rab9a in its GTP-bound form initially interferes with retromer association (11). Upon hydrolysis of GTP bound to Rab9a, L2 associates with GTP-Rab7 which recruits retromer. Hydrolysis of Rab7 then allows the dissociation of Rab7 and retromer (10, 11). This Rab9a/Rab7-mediated control of retromer association and dissociation is required for HPV trafficking from endosome to TGN. After arrival at the TGN, HPV L2 associates with GTP-Rab6a, which recruits BICD2 and dynein, thereby assembling a complex that mediates trafficking of HPV from the TGN (Fig. 6B).

Consistent with this sequential model, HPV-Rab6a association requires endosome acidification, functional γ-secretase and Rab7, all of which are required for HPV to exit the endosome. The preferential binding of both L2 and BICD2 to the GTP-bound form of Rab6a (Fig. 5E) (41–43) may facilitate the recruitment of BICD2 to L2. In addition to interacting with BICD2 and dynein, Rab6a may promote dynein-mediated HPV trafficking by recruiting dynactin, which forms a complex with dynein and activates dynein motility (42, 51), or by releasing LIS1 from a dynein idling complex (52).

We previously reported that BICD2 knockdown results in HPV accumulation in endosomes and the TGN (30). Thus, BICD2 functions in HPV transport from endosomes to the TGN, as well as for exit from the TGN. In contrast, Rab6a knockdown causes accumulation in TGN only. This raises a possibility that BICD2 and dynein are recruited to HPV in the endosome in a Rab6a-independent manner, possibly by as-yet-unidentified factor(s) such as another Rab protein. Consistent with this idea, HPV-dynein association at 8 hpi was indeed independent of Rab6a (Fig. 3A and 3B). However, once HPV enters the TGN, Rab6a is required to stabilize, maintain, or reinstate the HPV-BICD2 interaction to allow further dynein-mediated transport of the incoming virus. Because of the blockade to TGN exit, we cannot determine if Rab6a also acts at more downstream steps of entry, such as Golgi-to-ER trafficking, a step Rab6a can mediate for cellular cargo. It is likely that trafficking from the TGN and through the Golgi stacks involves iterative rounds of vesicle budding, transport, and fusion, and Rab6a may act repetitively during this process. Indeed, the reduced binding of GDP-Rab6a to L2 may allow the complex to dissociate upon GTP hydrolysis, only to reform at a later step when GTP-Rab6a is regenerated, providing an explanation for the apparent requirement for Rab6a cycling for HPV transport.

Although Rab6a is required for HPV-BICD2 association in infected cells, a C-terminal segment of L2 can directly interact with purified BICD2 *in vitro*, indicating that L2 can also bind BICD2 independently of Rab6a. Direct binding of BICD2 to L2 could play a role in assembling the complex in earlier entry steps, prior to the Rab6a-dependent step, and/or it could stabilize the complex between HPV, BICD2, and dynein. Our peptide pulldown studies show that Rab6a and BICD2 have overlapping but distinct sequence requirements for binding to L2. L2 434-461 peptide binds both Rab6a and BICD2, whereas peptide 442-461 binds BICD2 but not Rab6a, confirming that BICD2 can bind L2 independently of Rab6a. Retromer also binds to this segment of L2, so presumably the binding of various factors to this region of L2 is precisely orchestrated to allow proper trafficking. We have not yet identified L2 mutants defective for BICD2 or Rab6a binding that retain retromer binding, and retromer binding is required for stable L2 protrusion into the cytoplasm. Therefore, we cannot assess the consequences of L2 mutations that block BICD2 or Rab6a binding in infected cells because the available mutations prevent stable L2 protrusion.

Rab6a also plays important roles in infection by other viruses. Rab6a knockdown impairs herpes simplex virus type 1 (HSV1) infection by inhibiting HSV1 glycoprotein trafficking from Golgi to the plasma membrane and capsid envelopment (53). Thus, HSV1 requires Rab6 for anterograde trafficking during virion production, whereas HPV requires it for retrograde trafficking during entry. Rab6a may recruit kinesin as Rab6a interacts with kinesins and functions during anterograde trafficking (53, 54). In addition, Rab6 promotes production of infectious human cytomegalovirus, another herpesvirus, by supporting the trafficking of viral protein(s) into the viral assembly compartment (55). HIV infection also requires Rab6a, although the specific step Rab6a mediates is not known (56). Thus, different viruses appear to employ Rab6a for different functions during virus entry, assembly and/or, exit.

Our study elucidates a role of Rab6a in facilitating HPV entry by promoting HPV-BICD2- dynein association in the TGN, enabling retrograde trafficking of HPV from the TGN to the *cis*- Golgi. Rab6a engages with HPV by interacting with L2 after the incoming viruses has arrived at the TGN. This function of Rab6a not only sheds light on virus entry mechanisms but also provides insights into cellular protein trafficking, albeit with distinct Rab6a functions compared to known cellular cargo. Our findings also reveal sequential interactions between viral and host proteins during HPV entry, potentially identifying targets for therapeutic interventions to inhibit infection.

## Materials and Methods

### Cell lines

HeLa S3 cells were purchased from American Type Culture Collection (ATCC). HaCaT cells were purchased from AddexBio Technologies. 293TT cells were generated by introducing SV40 Large T antigen cDNA into HEK293T cells to increase Large T antigen expression and obtained from Christopher Buck (NIH). All cell lines were cultured at 37°C and 5% CO2 in Dulbecco’s modified Eagle’s medium (DMEM) supplemented with 20 mM HEPES, 10% fetal bovine serum (FBS), L-glutamine, and 100 units/mL penicillin streptomycin (DMEM10).

### Production of HPV pseudovirus (PsV)

HPV16 PsVs were produced by co-transfecting 293TT cells with wild-type p16SheLL- 3XFLAG tag (18) together with pCINeo-GFP [obtained from C. Buck] or pCINeo-Gluc (27) using polyethyleneimine (MilliporeSigma). For HPV18 and HPV5, p18SheLL-HA tag and p5SheLL were used, respectively. PsVs were purified by density gradient centrifugation in OptiPrep (MilliporeSigma) as described (57). Briefly, cells were washed with Dulbecco’s Phosphate Buffered Saline (DPBS) at 24 h post transfection, incubated in DMEM10, and collected 72h post transfection in siliconized tubes. The cells were then incubated in lysis buffer (DPBS with 0.5% Triton X-100, 10 mM MgCl2, 5 mM CaCl2, 100 µg/mL RNAse A (Qiagen)) overnight at 37°C in a water bath to allow capsid maturation. The lysates containing matured PsVs were loaded on an OptiPrep gradient that had been stabilized at least 1 h and centrifuged at 50k ×g for 3.5-4 h at 4°C in a SW-55Ti rotor (Beckman). Fractions were collected in siliconized tubes and subjected to SDS-PAGE followed by Coomassie blue staining to assess the abundance of L1 and L2 proteins. Peak fractions were pooled, aliquoted, and stored at -80°C.

### Determining HPV PsV infectivity

0.37x10^5^ HeLa S3 cells per well were plated in 24-well plates ∼48 h prior to infection. Approximately 6 h later, cells were transfected with 6.7 nM of indicated siRNAs (Table S1) using Lipofectamine RNAiMAX (Invitrogen) according to manufacturer’s protocol. Non- targeting siRNA (Table S1) was used as a negative control. 40-48 h after transfection, cells were infected with PsVs at an infectious MOI of ∼5 in unmodified HeLa cells. At 48 hpi, infectivity was determined by using flow cytometry to measure fluorescence intensity produced from expression of the reporter gene. Relative percent infectivity was determined by normalizing mean fluorescence intensity of samples transfected with experimental siRNA to that of the cells transfected with control siRNA, which was set at 100%.

### Western blot analysis

siRNA-treated cells were lysed using ice-cold 1X Radioimmunoprecipitation assay (RIPA) [50 mM Tris (pH 7.4), 150 mM NaCl, 1% Nonidet P-40, 1% sodium deoxycholate, 0.1% sodium dodecyl sulfate, 1 mM Ethylenediaminetetraacetic acid] buffer supplemented with 1X HALT protease inhibitor cocktail (Pierce) for 15 min at 4°C. After centrifugation at 14,000 rpm for 15 min at 4°C, the protein concentration in the supernatant was determined by Bicinchoninic acid (BCA) protein assay (Pierce). After normalization for protein amounts, the supernatant was mixed with 4X Laemmli dye (Bio-rad) supplemented with 10% 2-mercaptoethanol and incubated in a water bath for 7 min at 100°C. For pull-down experiment samples, please see below how samples were prepared. Samples were then separated by SDS-PAGE (4-12% gel) (Bio-rad) and analyzed by Western blotting using antibodies recognizing Rab6a (Thermo Fisher, 11420-1-AP, 1:1,000), BICD2 (Abcam, ab117818, 1:1,000), beta-actin (Sigma, A5441, 1:5,000), FLAG (Sigma, F1804, 1:1,000; Sigma, A8592, 1:1,000; Invitrogen, MA1-142, 1:1,000) or HA (Cell signaling, 3724, 1:1,000; Cell signaling, 2999, 1:1,000; Roche, 54732500, 1:1,000). Secondary horseradish peroxidase (HRP)-conjugated antisera recognizing rabbit, rat, or mouse antibodies as appropriate (Jackson ImmunoResearch, 711-035-152, 712-036-150, 115-035-146) were used at 1:5,000 dilution in 5% non-fat milk unless specified otherwise. The blots were developed with SuperSignal West Pico or Femto Chemiluminescent substrate (Pierce) and were visualized by using a film processor (Fujifilm).

### Proximity ligation assay (PLA)

0.35x10^5^ HeLa S3 cells per well were plated in 24-well plates containing glass coverslips 48 h prior to infection. Approximately 6 h later, cells were transfected with 6.7 nM of indicated siRNA (Table S1) as described above. At 40-48 h after transfection, cells were infected with PsVs at MOI of ∼200 in unmodified HeLa cells. As indicated, DMSO (0.2% as final concentration), 100 nM BafA, or 2 µM XXI were added to the medium 30 min prior to infection. At indicated times post-infection, cells were fixed with 4% paraformaldehyde (Electron Microscopy Sciences) at room temperature (RT) for 12 min, permeabilized with 1% Saponin (Sigma-Aldrich) at RT for 35-40 min, and blocked using DMEM10 at RT for 1-1.5 h. Cells were then incubated overnight at 4°C with a pair of mouse and rabbit antibodies: a mouse antibody recognizing L1 (BD Biosciences, 554171, 1:1,000 when used with anti-TGN46, 1:00 when used with other antibodies), FLAG (SIGMA, F1804, 1:1000), or HA (BioLegend, 901513, 1:200); and a rabbit antibody recognizing cellular proteins or epitope tags (anti-EEA1, Cell Signaling Technology, 2411, 1:75; anti-TGN46, Abcam, ab50595, 1:600; anti-GM130, Abcam, ab52649, 1:200; anti-dynein, Invitrogen, PA5-89505, 1:200; anti-FLAG, Cell Signaling Technology, 14793, 1:200). PLA was performed with Duolink reagents (Sigma Aldrich) according to the manufacturer’s instructions as described (58). Briefly, cells were incubated in a humidified chamber at 37°C with a pair of PLA antibody (mouse and rabbit) probes for 75 min, with ligation mixture for 45 min, and then with amplification mixture for 3 h, followed by series of washes.

Nuclei were stained with 4,6-diamidino-2-phenylindole (DAPI). Cellular fluorescence was imaged using the Zeiss LSM980 confocal microscope. Images were processed using a Zeiss Zen software version 3.1 and quantified using Image J Fiji version 2.3.0/1.53f.

### Protein purification

HEK 293TT cells were seeded in a 15-cm plate and transfected with plasmids expressing FLAG-BICD2, CA HA-Rab6a, or DN HA-Rab6a. After 48 hours, cells were washed three times with DPBS, harvested, and lysed in HN buffer (50 mM Hepes and 150 mM NaCl) with 1% triton and protease inhibitor cocktail (Thermo Fisher Scientific). Cells were incubated on a rotator for 20 min at 4°C then centrifuged for 15 min at 17,000 ×g. The resulting supernatant was incubated with anti-FLAG M2 or anti-HA agarose beads (Millipore Sigma, A2220; Pierce, 26181) for two hours at 4◦C. The beads were then washed once with the lysis buffer, twice with the lysis buffer containing 1 mM ATP to remove contaminating proteins, then once with HN buffer containing 0.1% Triton X-100 in HN buffer. All the buffers contain protease inhibitor cocktail. The proteins were then eluted twice with 3xFLAG or HA peptides in HN buffer containing 0.1% Triton X-100 at room temperature for 30 min. The quality of the purified proteins was analyzed by SDS-PAGE and SimplyBlue SafeStain (Invitrogen) and the quantity of them was determined using the BCA Protein Assay Kits (Pierce, 23225).

### Peptide pulldown

The peptides were purchased from GenScript or Promega and dissolved in DMSO. Stocks were diluted to 5 mg/mL and stored at -80◦C. Purified proteins (∼20 nM of FLAG-BICD2 or HA-tagged CA or DN Rab6a) were incubated with 5 µg of biotinylated peptide in HN buffer containing 0.1% triton, 1 mM DTT, and protease inhibitor cocktail (Thermo Fisher Scientific) for two hours at 4◦C with mild rotation. When specified, proteins were used at the indicated amounts. 30 µL of Pierce streptavidin magnetic beads (Thermo Fisher Scientific) were added to the samples and incubated for an additional hour at 4◦C. The beads were then washed three times with HN buffer and incubated at 95◦C for 10 min in SDS sample buffer with 2-mercaptoethanol. Precipitated proteins were analyzed by Western blot analysis.

### Co-immunoprecipitation of BICD2 and L2

HeLa cells were grown to ∼70% confluency in 10 cm plates before transfection and infection.

At 16 hpi, the cells were washed three times with DPBS. The second PBS wash contained 300 mM NaCl to remove extracellular HPV. Cells were lysed in 400 µL of RIPA buffer (50 mM Tris pH 7.4, 150 mM NaCl, 0.25% sodium deoxycholate, 1% NP40, and 1 mM EDTA) containing 1 mM PMSF. Cells were incubated on ice for 10 min followed by centrifugation at 16,100 ×g for 10 min. After centrifugation, 10% of the supernatants were taken for input and the remaining supernatants were incubated with BICD2 or the control IgG antibody overnight at 4◦C. Pierce protein A/G agarose beads (Thermo Fisher Scientific) were then added to the samples for 30 min at 4◦C, followed by three washes with the RIPA buffer and incubation at 95◦C for 10 min in 5x SDS sample buffer with 2-mercaptoethanol (i.e., elution). Samples were then analyzed by SDS-PAGE and Western blot analysis carried out as described above.

### Determining HPV PsV infectivity in cells transiently expressing CA or DN versions of Rab6a or Rab33b

0.35x105 293TT cells per plate were plated in 24-well plates 16-20 h prior to transfection of plasmids encoding HA-tagged CA or DN Rab6a or Rab33b, or the empty vector as control (Takara, 632105). 48 h after transfection, cells were infected with PsVs at MOI of ∼1 in unmodified HeLa cells. 48 hpi, cells were fixed with 4% paraformaldehyde, permeabilized with 1% Saponin, and blocked with 3% BSA. Cells were then stained using PE-conjugated antibodies recognizing HA (MACS Molecular, 130-120-717, 1:1000), followed by three to four times of washes using DPBS containing 0.1% Tween-20. HA-tagged Rab protein abundance and HPV PsV infectivity were determined by flow cytometry measuring fluorescence intensity produced by PE-labeled proteins and expressed GFP simultaneously.

### Immunofluorescence

HeLa cells were seeded onto glass coverslips in a 6-well plate and treated as indicated. Cells were washed three times with PBS, fixed in 4% paraformaldehyde for 20 min at room temperature then washed four times with PBS. Permeabilization was carried out for 20 min at room temperature with tris-buffered saline (TBS)/0.2% Triton X-100/3% BSA followed by three washes with TBS-T (TBS with 0.1% Tween-20). The cells were blocked with TBS (with 0.2% Tween-20 and 3% BSA) for one hour at room temperature. Primary antibodies (anti-BicD2, Novus Bio, NBP 2-43683, 1:200; anti-TGN46, Proteintech, 13573-1-AP, 1:2,000) were diluted in TBS (with 0.2% Tween-20 and 3% BSA) and incubated with the coverslips overnight at 4◦C. Secondary antibodies (goat anti-rabbit Alexa Fluor 488, Thermo Fisher Scientific, A11008, 1:2,000; goat anti-mouse Alexa Fluor 594, Thermo Fisher Scientific, A11032, 1:2,000) were diluted with TBS containing 0.2% Tween-20 and 3% BSA and incubated with coverslips for one hour at room temperature. Coverslips were mounted with mounting medium containing DAPI (Abcam, ab104139). Images were taken with confocal microscopy (Zess LSM 800 confocal laser scanning microscope with a Plan-Apochromat 40x/1.4 oil differential interference contrast M27 objective). Representative images were chosen out of three independent experiments.

### Construction of plasmids

Plasmids expressing HA-tagged Rab6a CA or DN variants were constructed as follows: CA and DN mutant Rab6a genes were amplified from EGFP-Rab6AQ72L (Addgene #49483) and EGFP-Rab6AT27N (Addgene, Plasmid #49484), respectively, using primers HA-Rab6a-BamHI-F (TGA ACC GTC AGA TCG CCT GGA GAA GGA TCC ATG TAC CCA TAC GAC GTT CCA GAT TAC GCT TCC ACG GGC GGA GAC TTC GGG AAT CCG) and Rab6a-EcoRI-R (GAA AAG CGC CTC CCC TAC CCG GTA GAA TTC TTA GCA GGA ACA GCC TCC TTC ACT GAC TGG TTG), then introduced between BamHI and EcoRI sites of pRetroX-Tight-Pur vector (Takara, 632105). Resulting plasmids containing the desired mutation were confirmed by DNA sequencing.

For protein purification purpose, pCMV-FLAG vector was used. Primers HAR6a-pCMV- XhoI-F (TAC AAG CTA CTT GTT CTT TTT GCA CTC GAG GCC ACC ATG ATG TAC CCA TAC GAT GTT CCA GAT TAC GCT TCC ACG GGC GGA G) and HAR6a-pCMV-AgeI-R (GTA TCT TAT CAT GTC TGC TCG AAG CGG ACC GGT TTA GCA GGA ACA GCC TCC TTC ACT GAC TGG TTG) were used to amplify CA or DN HA-tagged Rab6a, then introduced between XhoI and AgeI sites of pCMV-FLAG vector (Takara, 632105). Resulting plasmids containing the desired mutation were confirmed by DNA sequencing.

### Statistical analyses

For comparisons of two groups, unpaired *t*-tests were applied. For comparisons of more than three groups, One-way ANOVA with the ordinary ANOVA test. These analyses provide *P*- values for each comparison.

## Acknowledgments

We thank Qun Lin for making HPV18 and HPV5 PsVs. This work was supported by grants from the NIH to BT and DD (R01-AI150897) and to DD (R35-CA242462).

---

**Fig. S1.**
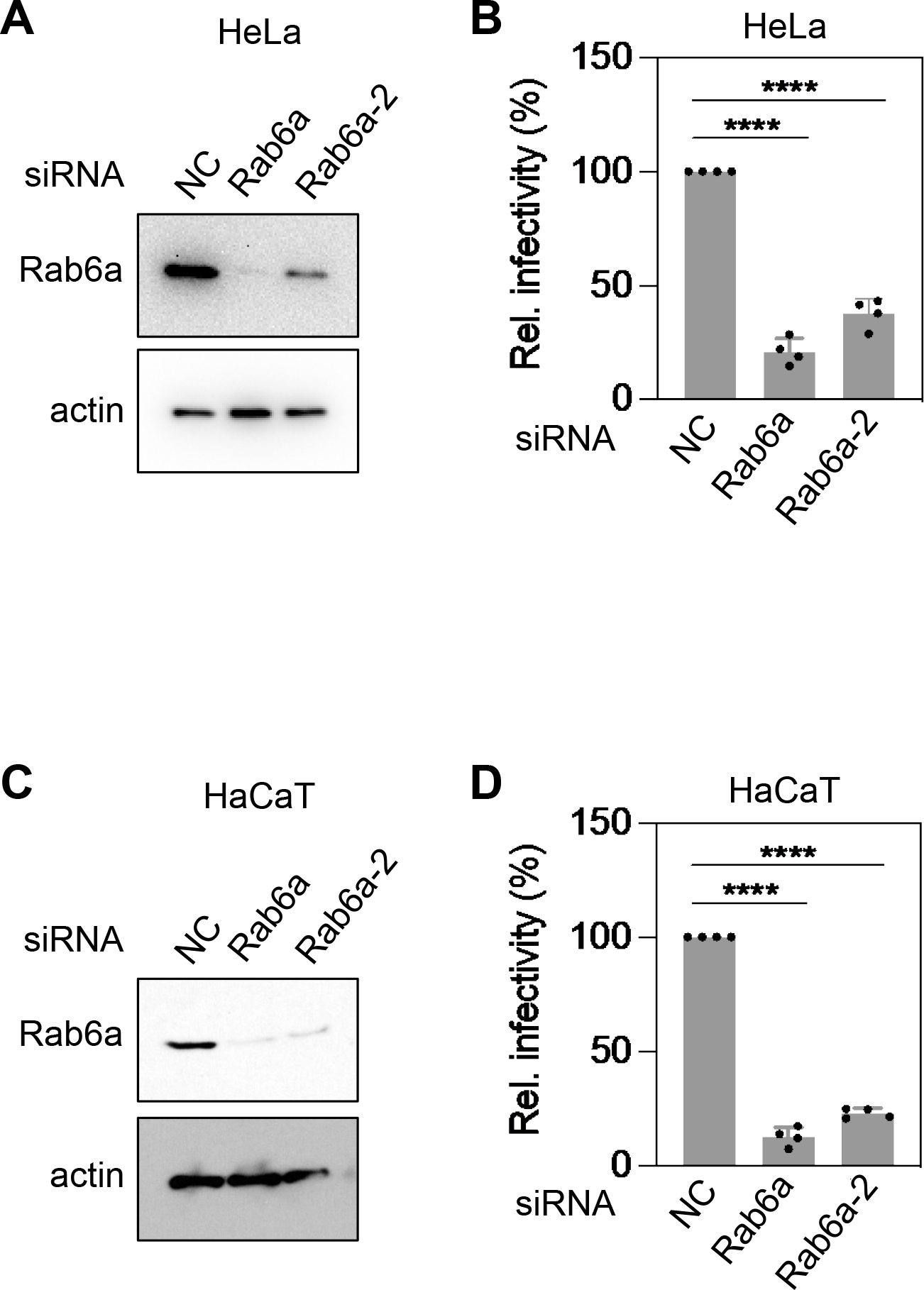
Rab6a is required for HPV entry in both HeLa and HaCaT cells. (**A**) HeLa S3 cells were transfected with siNC or two different siRNAs targeting Rab6a (siRab6a and siRab6a-2) and were subjected to Western blot analysis using antibodies recognizing Rab9a and actin as a loading control. (**B**) siRNA-treated cells as described in (**A**) were mock-infected or infected at the MOI of ∼2 with HPV16 PsV L2-3XFLAG containing the GFP reporter plasmid. At 48 hpi, GFP fluorescence was determined by flow cytometry. The results are shown as percent relative infectivity (based on mean fluorescence intensity) normalized to siNC treated cells (*right*). Each dot shows the result of an individual experiment. Bars and error bars show mean and standard deviation, respectively. ****, *P* < 0.0001. (**C**) As in (**A**) except using HaCaT cells. (**D**) As in (**B**) except using HaCaT cells.

**Fig. S2.**
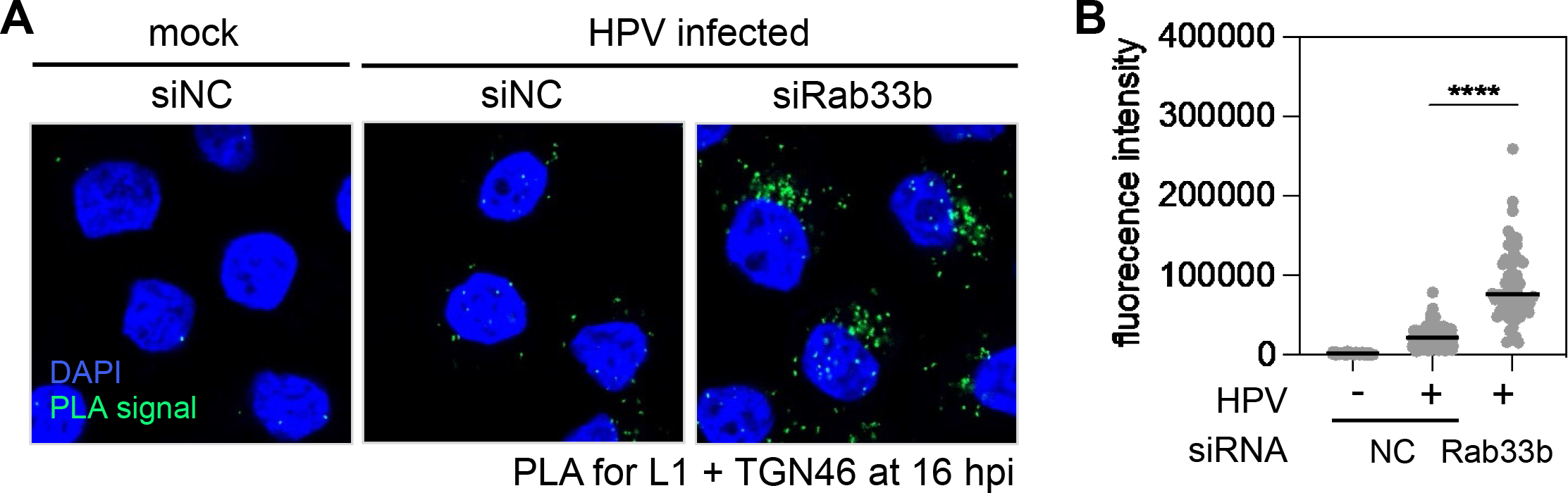
Excess of either GTP-Rab33b or GDP-Rab33b impairs HPV entry. (**A**) HeLa S3 cells were transfected with siNC or siRNAs targeting Rab33b and mock-infected or infected at the MOI of ∼200 with HPV16 PsV L2-3XFLAG containing the Gluc reporter plasmid. At 16 hpi, PLA was performed with antibodies recognizing HPV L1 and TGN46. Mock, uninfected; HPV, infected. PLA signals are green; nuclei are blue (DAPI). Similar results were obtained in two independent experiments. (**B**) The fluorescence of PLA signals was determined from multiple images obtained as in (**A**). Each dot represents an individual cell (*n*>40) and black horizontal lines indicate the mean value of the analyzed population in each group. ****, *P* < 0.0001. The graph shows results of a representative experiment.

**Table S1.** Oligonucleotides used in this study.

| **siRNA** | **source** | **identifier** |
| --- | --- | --- |
| siGENOME Human RAB6A siRNA<br>(shown as siRab6a in this study) | Dharmacon | D-008975-04-0005 |
| ON-TARGETplus Human RAB6A siRNA<br>(shown as siRab6a-2 in this study) | Dharmacon | J-008975-08-0002 |
| ON-TARGETplus Human RAB33B siRNA (SMARTPool)<br>(shown as siRab33b in this study) | Dharmacon | L-008909-00-0005 |
| ON-TARGETplus Human RGP1 siRNA (SMARTPool)<br>(shown as siRgp1 in this study) | Dharmacon | L-021128-02-0005 |
| ON-TARGETplus Human RIC1 siRNA (SMARTPool)<br>(shown as siRic1 in this study) | Dharmacon | L-026110-02-0005 |
| ON-TARGETplus Human Rab7A and Rab7B siRNA<br>(SMARTPool)<br>(shown as siRab7 in this study) | Dharmacon | L-010388-00-0005<br>L-018225-00-0005 |
| ON-TARGETplus Non-targeting Control Pool<br>(shown as siNC in this study) | Dharmacon | D-001810-10-20 |

